# Two-Dimensional Nonlinear Structured Illumination Microscopy with rsEGFP2

**DOI:** 10.1101/2025.05.11.653285

**Authors:** Shaoheng Li, Ryo Tamura, Kota Banzai, Daichi Kamiyama, Peter Kner

## Abstract

Superresolution microscopy enables imaging of subcellular structures and dynamics with nanoscale detail. Among the various superresolution techniques, structured illumination microscopy (SIM) stands out for its compatibility with live-cell imaging. Linear SIM is restricted to a resolution improvement of a factor of two, improving the resolution to about 100 nm. Nonlinear SIM (NSIM) utilizes reversibly switchable fluorescent proteins to generate a nonlinear response, allowing for the collection of higher spatial frequency information and theoretically extending the resolution without limit. By employing rsEGFP2 and patterned depletion illumination (PD) to generate the desired nonlinearity in the fluorescent response, we have successfully achieved 2D PD-NSIM imaging of actin in live U2OS cells with sub-80 nm resolution.

## 1. Introduction

Structured Illumination Microscopy (SIM), a popular superresolution method for live cell imaging, offers a unique approach to surpassing the diffraction limit from the perspective of the frequency domain [1, 2]. In SIM, the sample is illuminated with a sinusoidal pattern that modulates the emitted fluorescence intensity to induce Moiré fringes, lower-frequency patterns resulting from the mixing of the illumination pattern and the structure of the sample. These Moiré fringes are observable with the microscope even when the sample structure is not resolvable. Because the illumination pattern is known, the Moiré fringes can be used to reconstruct the sample structure beyond the resolution limit of the microscope.

One of the biggest advantages of SIM over other superresolution methods is the imaging speed. For live cell imaging, temporal resolution is important, and SIM has been demonstrated imaging whole live cells [3, 4]. Most of the current superresolution methods achieve remarkable spatial resolution often at the expense of diminished temporal resolution and/or diminished field of view. Since its invention [1, 2], SIM has been integrated with various imaging techniques including total internal reflection microscopy (TIRF) [3, 5, 6], light sheet microscopy [7, 8], and multifocal imaging [9–11]. However, the resolution enhancement of linear SIM is capped at a 2-fold improvement because the resolution improvement is proportional to the frequency of the structured illumination pattern. Consequently, achieving superresolution below 100 nm for *in vivo* live cell imaging with SIM remains challenging. This can be achieved with very high NA objectives that can use toxic immersion media [6].

The resolution of SIM can be extended by harnessing the nonlinear response of fluorophores. When the fluorescence emission is a nonlinear function of the excitation intensity, higher harmonics of the structured illumination pattern are generated, resulting in higher-frequency information being shifted into the passband of the microscope. As the nonlinearity increases, more high frequency harmonics emerge, enabling higher spatial frequencies to be mixed down. Theoretically, the extent of the resolution improvement is unlimited since it is dictated by the number of high-frequency harmonic terms [12]. In reality, however, the final resolution is affected by the noise in the system, and only the frequency terms above the noise level contribute to the resolution of the final image [13]. Extending the resolution of SIM with the nonlinear response of the fluorophores is referred to as Nonlinear SIM (NSIM).

There are different ways of generating the desired nonlinearity. NSIM was first demonstrated using excitation saturation as the nonlinearity [12]. But excitation saturation requires very high excitation intensities and is not compatible with live-cell imaging. Reversibly switchable fluorescent proteins (rsFPs) that can toggle between a fluorescent and a non-fluorescent state can generate a non-linear fluorescent response at much lower excitation intensities and have been used for most NSIM realizations. rsFPs typically use two wavelengths. Negatively photoswitchable rsFPs (np-rsFPs) use one wavelength for activation and a longer wavelength for both excitation and deactivation. Positively photoswitchable rsFPs use one wavelength for activation and excitation and another wavelength for deactivation. NSIM with rsFPs was first demonstrated with the np-rsFP Dronpa [14]. NSIM has since been developed using several different np-rsFPs. NSIM has also been used with the positively photoswitchable rsFP (pp-rsFP) Kohinoor [15].

In principle the activation, deactivation, and excitation light can all be structured. However, for a np-rsFP, the activation and deactivation/excitation beams are at different wavelengths and care must be taken to match the patterns accounting for chromatic aberrations. To avoid this problem, patterned depletion NSIM (PD-NSIM) can be used. In this approach, for an np-rsFP, all fluorophores are activated with uniform illumination at the activation wavelength. Then the sample is illuminated with a depletion pattern that turns off the rsFPs, and, finally, the sample is illuminated with an excitation pattern that is *π* out of phase with the depletion pattern. This approach requires 3 separate illuminations per raw image.

To reduce the number of required illuminations, patterned activation NSIM (PA-NSIM) has been used [6]. In PA-NSIM, the sample is illuminated with a sinusoidal activation pattern and then illuminated with a sinusoidal excitation pattern. So only two exposures are needed rather than three. However, care must be taken to ensure that the activation and excitation patterns are matched since they are at different wavelengths and chromatic aberrations may affect their relative magnification. As the nonlinearity is increased in PA-NSIM, the background increases because the DC term increases with increasing nonlinearity.

So far, the rsFPs that have been used for NSIM include Dronpa [14], Skylan-NS [6, 16], rsEGFP2 [16], and Kohinoor [17]. Rego et al. succesfully demonstrated a lateral resolution of 40 nm using PD-NSIM with Dronpa to image purified microtubules, and nuclear pores in the intact nuclear membrane, capturing 63 raw readout images (7 phases and 9 orientations) [14]. They also imaged actin in intact fixed cells at 60 nm resolution. The switching rate of Dronpa is slow, requiring about 0.4 seconds of exposure to turn off with approximately 20 W/cm^2^ at 488 nm. This limits the imaging speed. Another significant limitation with Dronpa was its inability to go beyond 15 switching cycles before substantial bleaching set in. Achieving even this limited performance necessitated the use of toxic anti-bleaching chemicals, which limits its application in live-cell NSIM imaging. Li and and colleagues utilized Skylan-NS to perform PA-NSIM, imaging intracellular dynamics in living cells. They claimed a spatial resolution of approximately 60 nm and a time resolution of around 40 frames per second. Compared with other rsFPs, the contrast ratio of Skylan-NS reduces as the number of switching cycles increases, reducing the nonlinearity [16]. All resolution values reported by Li are stated as “theoretical”, calculated based on the number of orders used in NSIM, and the resolution is not independently evaluated [18]. Zhang et al. further compared Skylan-NS with rsEGFP2 and Dronpa using PA-NSIM and claimed that Skylan-NS outperforms other rsFPs for PA-NSIM [16]. While Skylan-NS offers higher photon yields and contrast ratios, rsEGFP’s faster switching kinetics, longer fluoresence decay, and low light intensity requirements also give it advantages. Although the studies [16, 19] show that rsEGFP2 is not ideal for PA-NSIM, its potential for PD-NSIM hasn’t been well studied.

Here, we present the first demonstration of live 2D PD-NSIM using rsEGFP2. We demonstrate imaging of actin filaments in U2OS cells, achieving sub-80 nm resolution in live-cell imaging. We examine the images in both in both real-space and Fourier-space to support our claim of increased resolution.

## 2. Methods

### 2.1. Principle of PD-NSIM

In fluorescence microscopy, each image is the product of sample structure and the excitation pattern convolved with the microscope point spread function (PSF), as shown in Fig. 1(a). If the excitation pattern is sinusoidal, it will create additional copies of the sample structure that are shifted in spatial frequency. In frequency-space additional band-limited copies of the sample information are generated as shown in Fig. 1(b). With SIM, the excitation pattern cannot have a sample frequency greater than 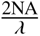, and the copies cannot be shifted further than the green circles shown in Fig. 1(b). Thus a frequency outside that region will still be unresolvable. In NSIM, the nonlinearity distorts the fluorescence emission pattern generating higher harmonics which create additional copies of the sample information shifted by larger amounts as shown in Fig. 1(c). The pattern is rotated to cover an isotropic region of frequency-space, Fig. 1(d). Excellent explanations of linear SIM can be found in [20].

**Fig. 1.**
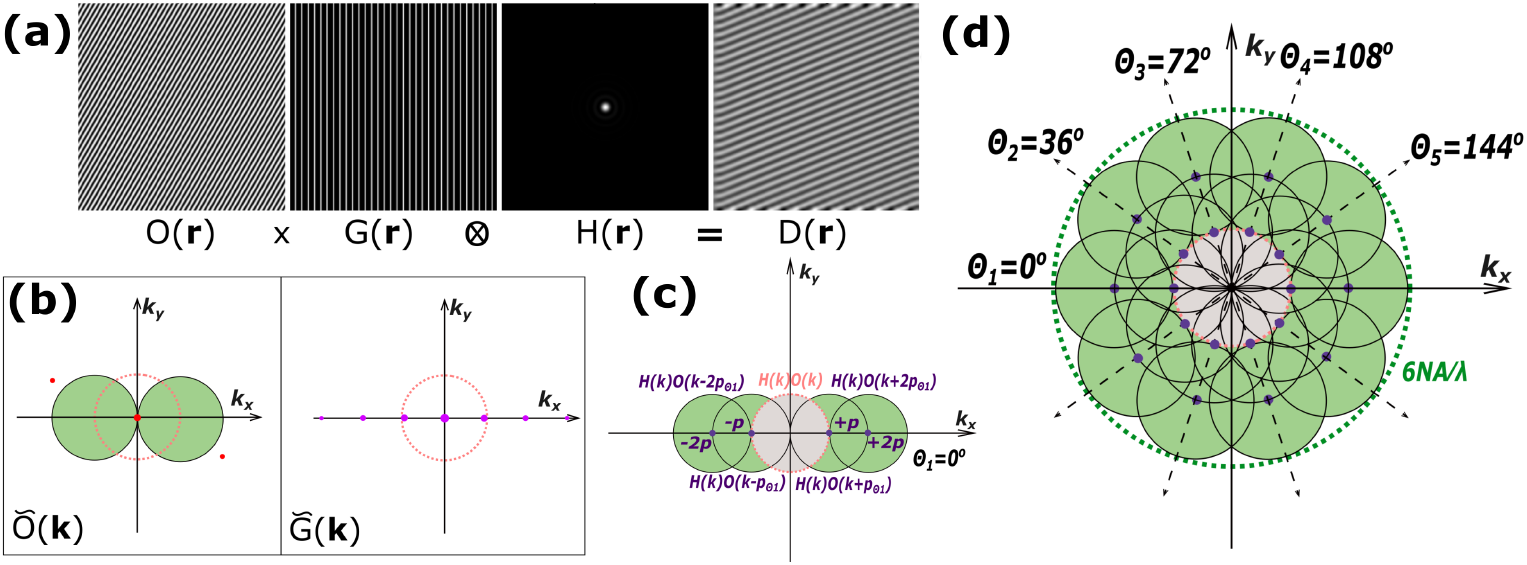
(a) Imaging formation of NSIM. (b) Frequency-space representation of object and effective illumination pattern. (c) Frequency content of first higher-order harmonic order NSIM within 5 circular regions. (d) Effective OTF of first higher-order harmonic order NSIM after rotating 5 angles.

In patterned depletion NSIM, the sample is illuminated three times for every exposure. First a uniform beam at the activation wavelength, 405 nm, activates the sample. Then the sample is illuminated with a patterned depletion beam at the depletion/excitation wavelength, 488 nm. This beam serves to deactivate the fluorophores. The longer the beam is left on, the more the fluorophores are deactivated. However, at the intensity zeros of the pattern, the fluorophores will not be deactivated. So, the longer the depletion is turned on, the narrower the peaks of activated fluorophores as shown in Fig. 2. The fluorescence is then excited with a third beam that is *π* out of phase with the depletion beam so that the intensity maxima are now centered on the remaining activated fluorophores. The excitation is then more sharply peaked than a sinusoid resulting in higher harmonics that provide higher resolution information.

**Fig. 2.**
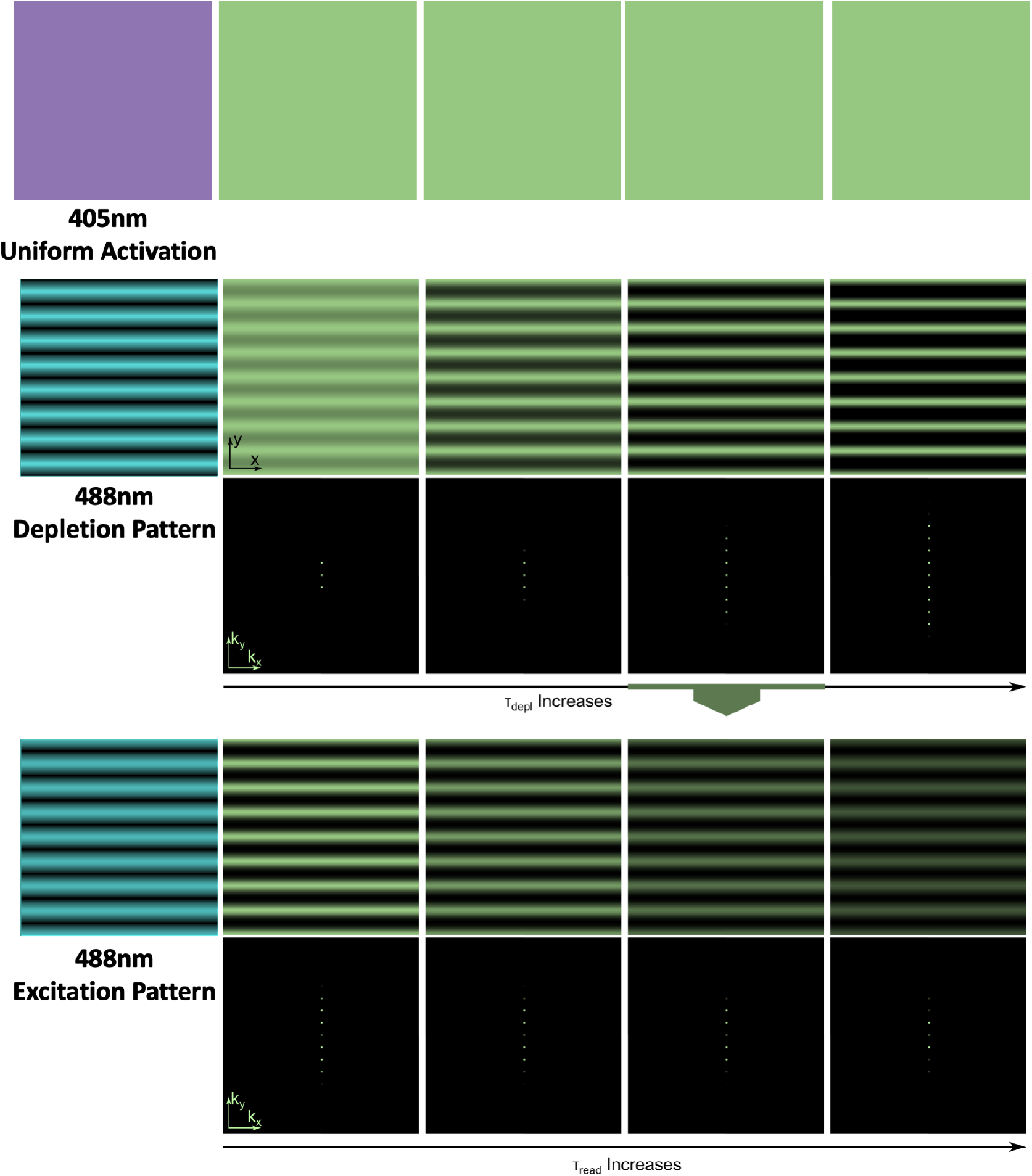
Patterned depletion Illumination for NSIM

### 2.2. Characterization of rsEGFP2

rsEGFP2 is an rsFP derived from the enhanced green fluorescent protein (EGFP) by replacing threonine 65 by alanine [21]. Fig. S1(a) shows the 3D structure of the protein. It operates in a negative switching mode, with a 480 nm depletion/excitation wavelength, and a 405 nm activation wavelength. rsEGFP2 is significantly faster than rsEGFP. At a light intensity of 5.5 kW/cm^2^, rsEGFP2 switches off about 6.5 times faster than rsEGFP. With a light intensity of 100 W/cm^2^, rsEGFP2 can achieve an off-rate greater than 90% with a 4ms exposure time at 480 nm. Similarly, rsEGFP2 achieves greater than 90% activation with a 5ms exposure to 250 W/cm^2^ at 405 nm. rsEGFP2 can undergo over 2,100 switching cycles without a significant reduction in fluorescence intensity, making it highly durable for PD-NSIM which requires more exposures than PA-NSIM. The relatively low light intensities required for switching between states help reduce phototoxicity and photobleaching making rsEGFP2 well suited for long-term live-cell imaging. Its successful application in RESOLFT further demonstrates its utility in high-resolution imaging [21].

Compared to Skylan-NS, the switching time of rsEGFP2 is 4 ms compared to 10 ms for Skylan-NS [16]. rsEGFP2 is a factor of 4 less bright than Skylan-NS. rsEGFP2 is more photostable than Skylan-NS, losing 20% of intensity over 1000 switching cycles and having a stable contrast ratio of 20. Whereas, Skylan-NS loses over 50% of intensity and the contrast ratio degrades from 37 to 17.

### 2.3. Optical Setup

Fig. 3 shows our experimental setup for 2D PD-NSIM. The system is built on an Olympus IX71 inverted microsope with a Prior Proscan XY Stage and a Prior Nanoscan SP200 Z-stage. The system includes a 405 nm activation laser (Coherent OBIS 405 nm LX 100 mW) and a 488 nm depletion/excitation laser (Coherent OBIS 488 nm LX 150 mW). The depletion beam is reflected by a polarizing beam splitter (PBS, 10FC16PB.3 Newport), through an achromatic half-wave plate (AHWP10M-600, Thorlabs) onto a spatial light modulator (SLM, Forth Dimension QXGA-3DM). The combination of PBS, HWP, and SLM generates binary phase-patterns as described in [3]. The activation beam is reflected by the PBS into the excitation path without reflecting off of the SLM, providing a uniform illumination to activate all the rsFPs without patterning. The activation and depletion beams are sent through a series of relay lenses to the microscope body which contains the tube lens and objective (UPLAPO60XOHR TIRF Objective, 60x, 1.50 NA, Olympus). The total magnification from the SLM to the sample is 240×. The relay lenses allow for the alignment of the illumination beams onto the sample with mirrors M5 and M7 which are close to the sample and pupil conjugate planes. Relay lenses L7 and L8 are used to create an image of the pupil at mirror M9 which can be replaced by a deformable mirror for adaptive optics correction. For this work, M9 is a dielectric mirror.

**Fig. 3.**
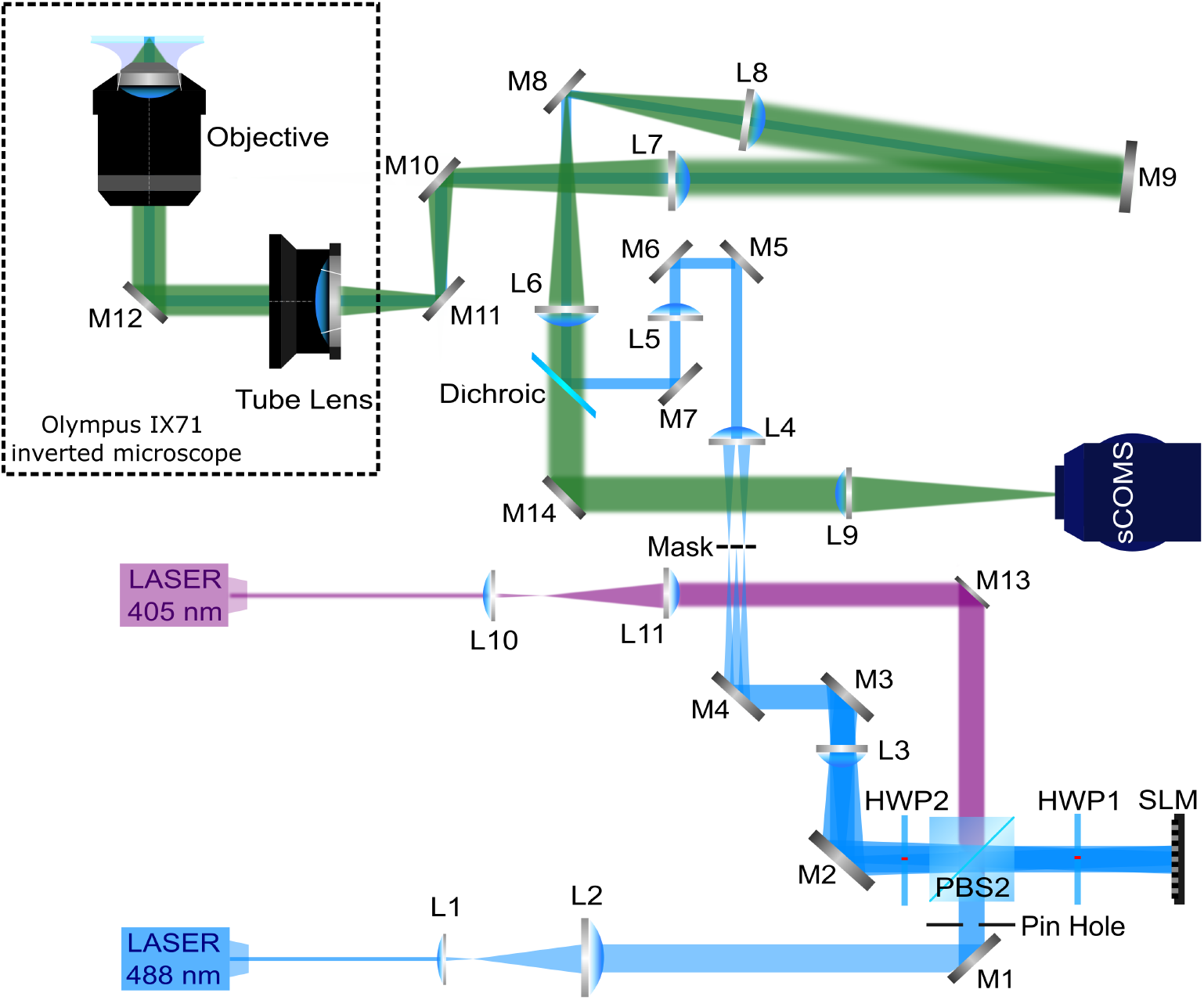
Experimental setup of 2D-NSIM. HWP: half-wave plate. PBS: polarized beam splitter. PH: pinhole. M1-M4: flat mirror. L1-L11: lens. *f*_1_ = 50 mm, *f*_2_ = 400 mm, *f*_3_ = 400 mm, *f*_4_ = 125 mm, *f*_5_ = 125mm, *f*_6_ = 100mm, *f*_7_ = 250mm, *f*_8_ = 250mm, *f*_9_= 300mm, *f*_1_0 = 50mm, *f*_1_1 = 200mm. The blue represents the illumination path (488nm) and the green path is the detection path.

The fluorescence emission is directed to the sCMOS camera (Andor SONA 4BV6U, 6.5*μ*m pixel size) through the “image flat” long-pass dichroic mirror (Di03-R488-t3-25×36, Semrock). The total magnification from the sample to the camera is 180×, and the effective pixel size is 36.1 nm. An emission filter (Semrock BrightLine quad band bandpass filter, FF01-446/523/600/677-25) and a notch filter (Semrock StopLine quad-notch filter, NF03-405/488/561/635E-25) are put before the camera to block unwanted light, assuring a low background noise level.

To maximize the modulation strength of the sinusoidal pattern, the polarization state of the two interference beams must be normal to the direction of the pattern wave vector (s-polarized) to achieve maximum contrast of the pattern. Therefore, the polarization state of the incident beam needs to be rotated along with the pattern orientation. A half-wave plate (HWP2 in Fig. 3) mounted in a fast-motorized rotation stage (8MRU, Altechna) is used to control the polarization. All but the ± 1 diffraction orders are filtered out by the mask after mirror M4 (custom design, Chrome mask on Soda Lime substrate, Photosciences, Inc.). The activation beam is tilted to align with one of the first-order diffracted beams of the depletion beam to ensure that it will pass through the mask.

### 2.4. Data Acquisition

The data acquisition process for 2D PD-NSIM begins by activating all fluorophores using uniform illumination of ∼ 10 W/cm^2^ for ∼ 5 ms at 405 nm. Then the patterned depletion illumination of 5 W/cm^2^ for 10 ms at 488 nm is applied. Finally, the phase of the illumination pattern is shifted by *π*, and the 488 nm excitation beam is turned on for 6 ms at ∼ 5 W/cm^2^.

The pattern at the SLM has a period of 15 pixels split 7/8 between positive and negative phase pixels. The nonlinear pattern generation is repeated with 5 phase shifts and 6 angles of rotation. Patterns at angles 0°, 30°, 60°, 90°, 120°, and 150° were used. A total of 30 raw images is required to reconstruct a single 2D PD-NSIM image. To increase SNR while maximizing the nonlinear response of rsEGFP2, we repeat the acquisition process 5 times for each phase, rather than using a single long exposure. The images are then averaged to create a final image stack for reconstruction.

### 2.5. Cell Culture and Transfection

The U2OS cells used in this paper were cultured in Dulbecco’s Modified Eagle Medium (DMEM) supplemented with 10% (vol/vol) fetal bovine serum (FBS) and 1% (vol/vol) penicillin-streptomycin (Pen-Strep) and maintained at 37°*C* and 5% (vol/vol) CO_2_ in a humidified incubator. Cells were passaged upon reaching apporximately 80-90 % confluence by washing with phosphate-buffered saline (PBS), detaching with 500*μ*L of trypsin-EDTA solution, neutralizing with four volumes of complete medium and centrifuging at 1500 rpm for 5 minutes. The pellet was resuspended in fresh growth medium and seeded at an appropriate density for subsequent use.

To visualize actin structures, the Actin-chromobody plasmid (a gift from Testa’s lab) was obtained. We performed standard transformation into E. coli strain *α*5, followed by inoculation into a large flask (50 mL culture volume) and overnight culture at 37° C. The next day, after centrifugation, the bacterial pellet was collected, and the plasmid was purified using a Qiagen midiprep kit. The purified plasmid, dissolved in ddH_2_O at a concentration of 1 − 2µg/µL, was used for subsequent transfection into U2OS cells.

The transfection was performed using PEI reagent. The imaged U2OS cells were plated on 35 mm ibidi dishes pre-coated with 0.1% PBS-diluted fibronectin human plasma and incubated overnight to achieve 90-95% confluence. For transfection, 3*μ*g of rsEGFP2 plasmid DNA was diluted in 50*μ*L of Opti-MEM medium, while 6*μ*L of PEI reagent was diluted in another 50*μ*L of Opti-MEM. The diluted PEI was added to the DNA solution, mixed thoroughly, and incubated at room temperature for 10 minutes to form complexes. The transfection mixture was added dropwise to the cells and incubated at 37°C and 5% CO_2_ for 24 hours. The cells were then imaged within 6 hours.

## 3. Results

### 3.1. Modulation Strength

Fig. S2 shows simulation results demonstrating how depletion and excitation exposures affect modulation strength. The modulation strength is measured as the OTF overlap magnitude. As expected, with excitation expsoure time fixed at 10 ms, an increase in depletion exposure time enhances modulation strength for both linear and nonlinear orders. Conversely, with depletion exposure fixed at 10 ms, longer excitation times diminish modulation strength for both harmonic order. However, the simulation results only show the effects of the depletion and excitation illumination exposures and do not account for specific characterists of rsEGFP2 such as the switching-off time, activation time, or photobleaching. We did measurements using rsEGFP2 labeled actin in U2OS cells; the results are shown in Fig. 4. We measure the modulation contrast for both the first and second order signals as a function of the excitation exposure or depletion exposure time. Each measurement was repeated on five different cells, and we plot the error bars from the different measurements.

**Fig. 4.**
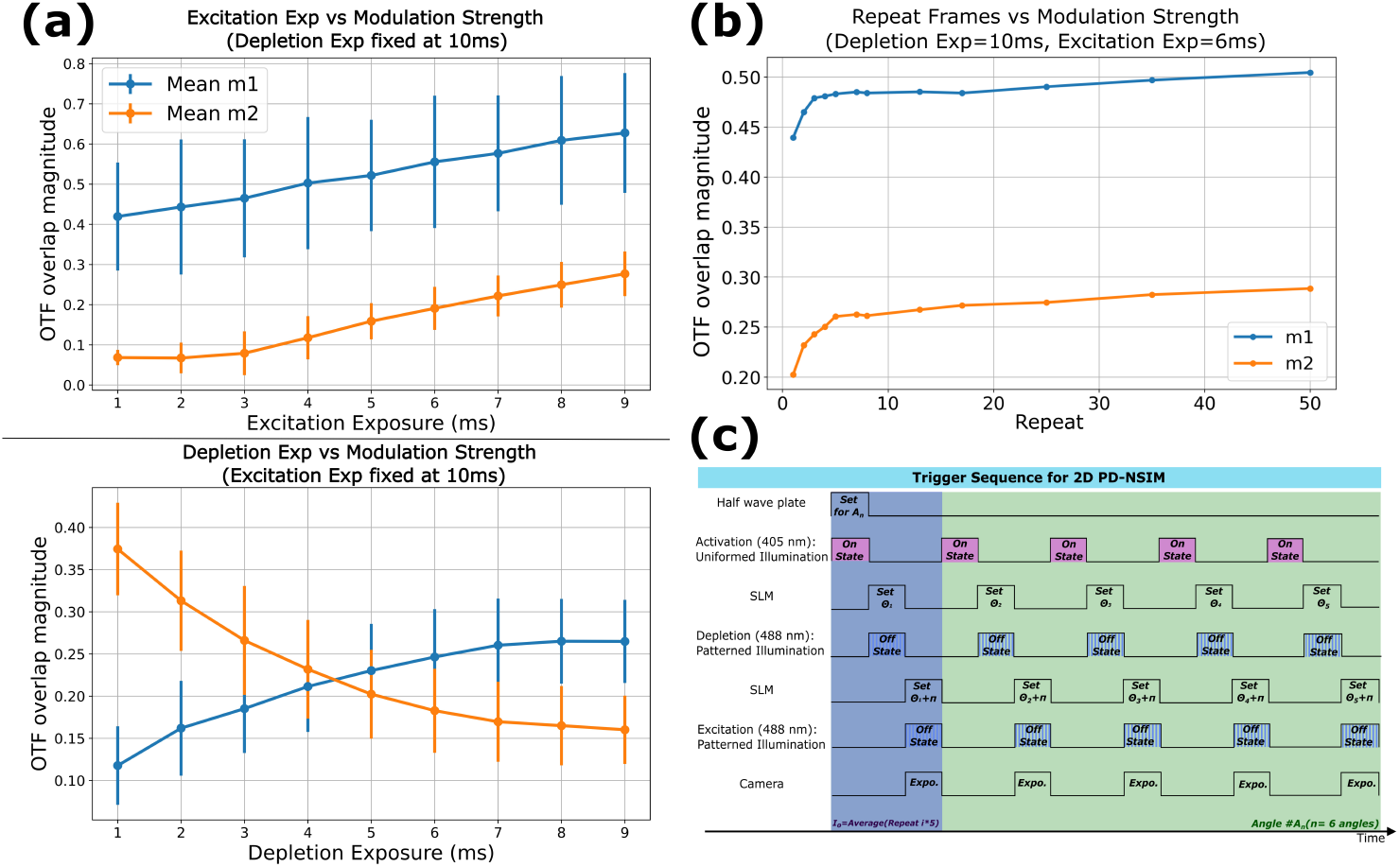
(a) Experimental results showing the effects of depletion and excitation pattern exposure on the nonlinear response. Data was collected from rsEGFP2-labeled U2OS cells with a coarse pattern spacing of 0.5NA / *λ*. For each plot, five cells were imaged, and the data represents the mean values with error bars indicating the standard deviation. Top inset: Relationship between excitation exposure time and modulation depth, with depletion exposure time fixed at 10 ms. Bottom inset: Relationship between depletion exposure time and modulation depth, with excitation exposure time fixed at 10 ms. (b) Experimental result of repeating test. Data was collected from same U2OS cell with an illumination pattern spacing of 0.5NA / *λ*. (c) Synchronization trigger sequence of all components during 2D PD-NSIM data acquisition for a single imaging cycle at one angle.

For excitation exposure, similar to the simulation results, increasing exposure time led to higher modulation strength for both harmonic orders. However, in the case of depletion, the modulation contrast for the first harmonic initially increased with longer depletion times, leveling off at 0.25 at 6 ms. In contrast, for the second harmonic, increasing depletion exposure resulted in a reduction in modulation strength. This may be due to a lower than desired modulation contrast of the excitation pattern. This will result in the fluorophores being turned off everywhere if the depletion beam is kept on too long. Therefore, to balance the linear and nonlinear response, we chose 10 ms for the depletion exposure and 6 ms for the excitation exposure.

To increase the SNR of the collected data, we repeated each imaging cycle multiple times and averaged the images. To determine the number of times to repeat each image, the illumination cycle was repeated with the setting of 10 ms depletion exposure and 6 ms excitation exposure for up to 50 cycles. The results, Fig. 4(c), show the modulation strength increases rapidly initially but only very slowly after 5 cycles. The increase in OTF overlap indicates an increase in the SNR. Therefore, we chose to repeat each raw image cycle 5 times. Fig. 4(d) shows the timing sequence of the lasers, polarization rotation, SLM, and camera.

### 3.2. NSIM

The performance of 2D PD-NSIM was demonstrated by imaging actin labeled live U2OS cells transfected with rsEGFP2. Results are shown in Fig. 5, Fig. S4, and Fig. S5. Fig. 5(a) shows live cell imaging of the U2OS cells with a pattern frequency of 1.66NA / *λ*. The effective OTF after the NSIM reconstruction has an extended bandwidth of 5.32NA / *λ*, as shown Fig. 5(b), 2.66 × the extent of the widefield OTF. This corresponds to a resolution of ∼ 70 nm. A clear improvement in resolution can be observed between WF, SIM, and PD-NSIM images as shown in the insets in Fig. 5(b) and (c). The Fourier Transform (FFT) of the spatial images also reveals strong features in frequency space, demonstrating the extended frequency space captured by PD-NSIM. Fig. 5(d) shows a different cell imaged with the same pattern spacing. A line profile through an actin fiber is shown in Fig. 5(e). The measured Full-Width at Half-Maximum (FWHM) of the fiber measured from the NSIM image is measured to be 80 nm, three times smaller than the measurement from the WF image. This improvement is greater than the increase in the OTF extent, because the Wiener filtering provides a further increase in the measured resolution.

**Fig. 5.**
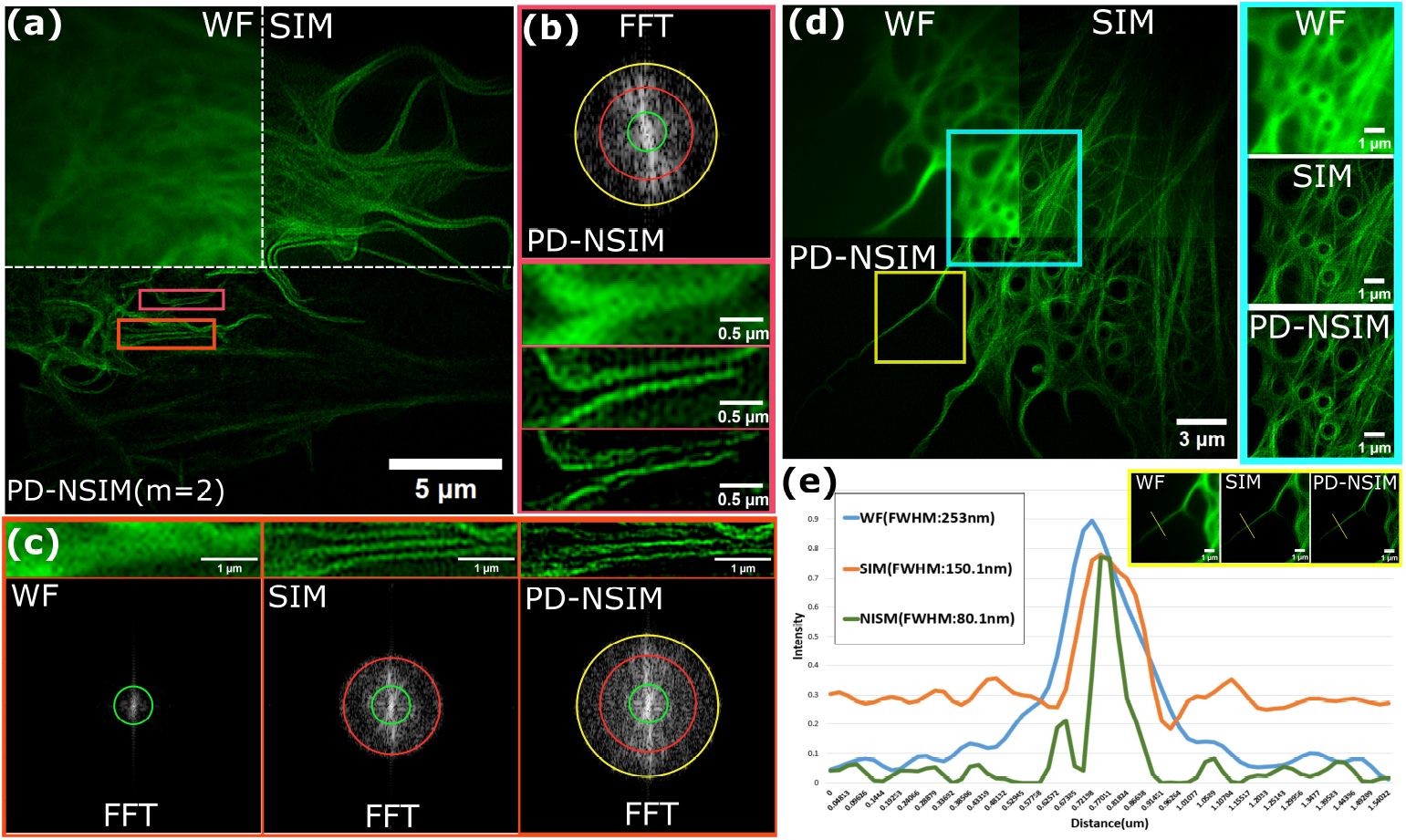
Live-cell imaging of 2D PD-NSIM with rsEGFP2. (a) A live U2OS cell transfected with rsEGFP2 imaged with widefield (WF), Linear SIM, and 2D PD-NSIM. (b) Bottom inset: zoomed-in view of the pink boxed region in (a). Top inset: corresponding fast Fourier transform (FFT) image. (c) Top inset: zoomed-in view of the orange boxed region in (a). Bottom inset: corresponding fast Fourier transform (FFT) image. (d) A live U2OS cell transfected with rsEGFP2 imaged with WF, Linear SIM, and 2D PD-NSIM. Right inset: Zoomed-in view of the blue boxed region in the left construction image. (e) Intensity plot of the yellow line marked in yellow boxed region from (d). The FFT image is displayed with logarithmic scaling for clarity with the green, red, and yellow circles represent effective OTF for WF, Linear SIM and the NSIM reconstructions in frequency domain.

Theoretically, for a pattern spacing of 1.66NA / *λ* (NA=1.5 and *λ* = 503 nm), the WF resolution is 167.6 nm. From simulations, the resolution after SIM and NSIM reconstruction with a Wiener filter is 92 nm and 68 nm respectively. To better demonstrate the repeatability of our approach and the real resolution improvement, we quantified the FWHM of 19 actin filaments from 7 different U2OS cells. We measured the filaments about 1*μ*m from the edge of the actin fiber to increase the chance that we are measuring a single actin fiber, and the FWHM represents the system resolution. Fig. S3 shows the resolution distribution of the measured actin fibers, giving a mean FWHM of 208 nm for WF, 97 nm for SIM, and 70 nm for PD-NSIM. The resolution improvement of PD-NSIM is 3×.

## 4. Conclusion

We successfully achieved live-cell superresolution imaging using 2D PD-NSIM with rsEGFP2. We optimized the data acquisition based on the characteristics of rsEGFP2 to enhance both the nonlinearity and modulation contrast. Live-cell imaging was successfully demonstrated with sub-80 nm resolution. In the future we plan to use PD-NSIM to perform time-lapse imaging of actin dynamics.

Several promising developments could further enhance the utility and performance of 2D PD-NSIM. One promising direction is to combine PD-NSIM with TIRF or light-sheet illumination which could improve the SNR and modulation contrast, and thereby increase the strength of the nonlinear harmonic orders [14, 19]. Another potential direction is extending 2D PD-NSIM into three-dimensions. By implementing 3D structured illumination patterns and performing axial scanning superresolution volumetric live-cell imaging could be achieved. Additionally, integrating Adaptive Optics into PD-NSIM could significantly improve image quality, and may even be essential for 3D PD-NSIM, particularly for deep tissue imaging, where optical aberrations pose substantial challenges [22, 23].

## Acknowledgments

We thank Dr. Ilaria Testa for providing the actin-chromobody plasmid.

