## Supplemental Text and Figures for "Two-Dimensional Nonlinear Structured Illumination Microscopy with rsEGFP2"

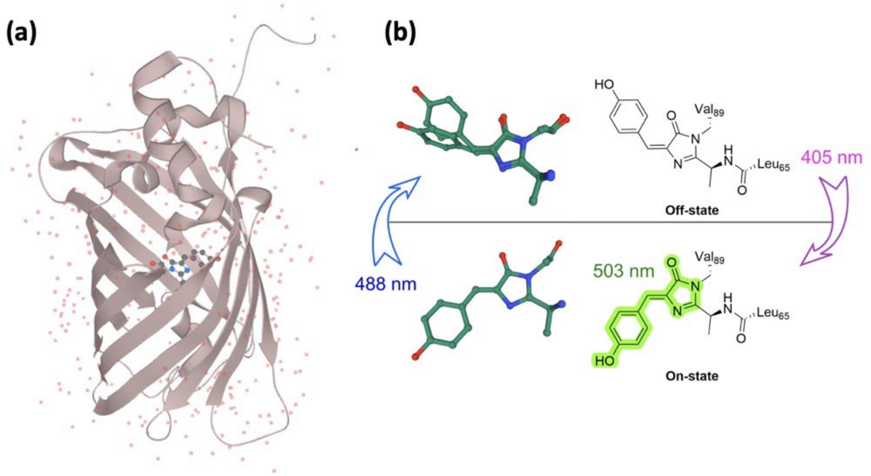


Fig. S1. (a)Three-dimensional structure of rsEGFP2 (PDB: 5O89). (b)The chromophore of rsEGFP2. In the ON state, it absorbs 488 nm light and emits at 503 nm, while isomerization shifts it to the OFF state. In the OFF state, 405 nm light triggers isomerization back to the ON state [1].

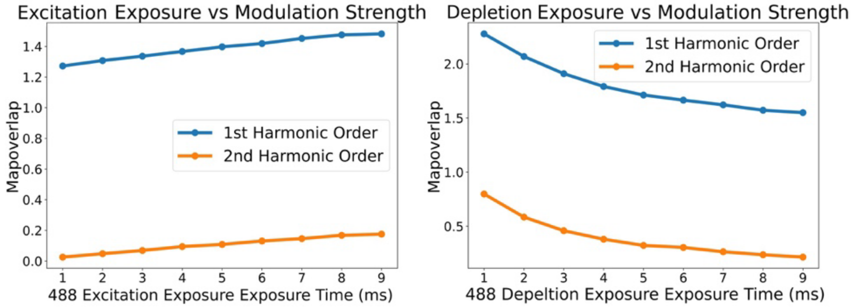


Fig. S2. Simulation results showing the effects of depletion and excitation pattern exposure on the nonlinear response. The pattern frequency is set near the diffraction limit at of2N A/λ. (a) Relationship between depletion exposure time and modulation strength, with excitation exposure time fixed at 10 ms. (b) Relationship between excitation exposure time and modulation strength, with depletion exposure time fixed at 10 ms.

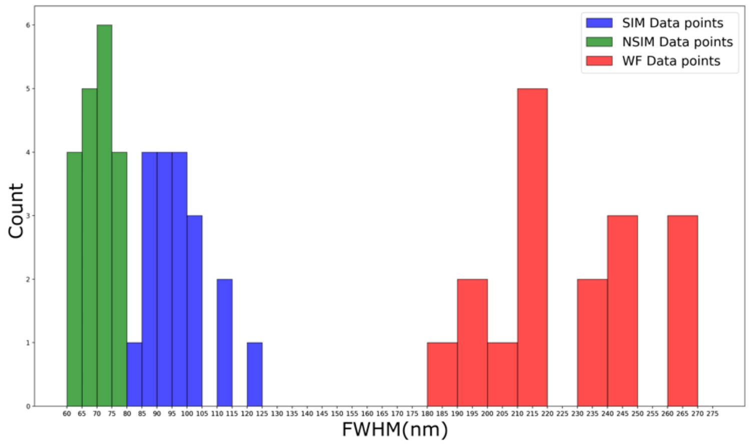


Fig. S3. Resolution distribution of the measured 18 U2OS actin fibers for WF, SIM, and NSIM. The resolution was measured by the FWHM of actin fibers.

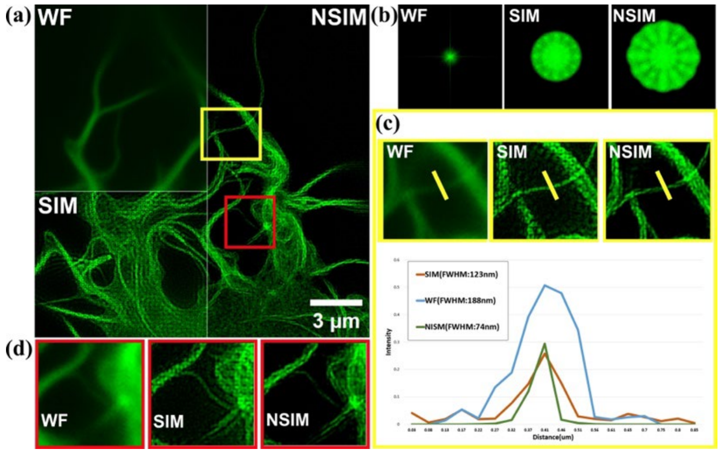


Fig.S4. (a) A fixed U2OS cell transfected with rsEGFP2 imaged with PD-NSIM. (b) Corresponding frequency distribution of the imaged cell in Fourier space. (c) Top figures: Zoomed-in view of the yellow boxed region in (a). Bottom figures: Intensity plot of the yellow line marked in top figures. The full width half maximum (FWHM) is measured for corresponding images. (d) Zoomed- in view of the red boxed region in (a).

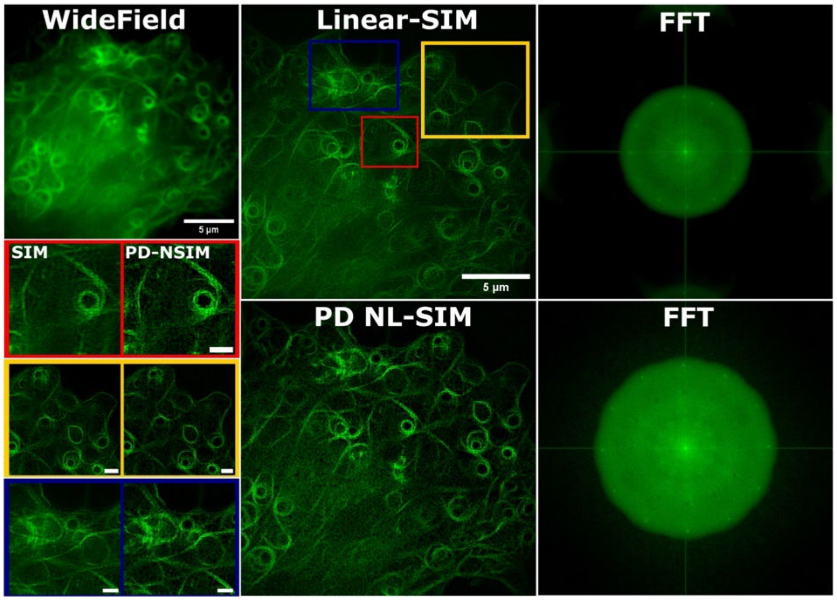


Fig. S5. Alive U2OS cell transfected with rsEGFP2 imaged with widefield, Linear SIM, and 2D PD- NSIM. Bottom left figures: Zoomed-in view of the yellow, red, and blue boxed regions in linear-SIM construction image, the scale bar indicated 1um. The FFT image is displayed with logarithmic scaling.
